# Distinct recruitment of feed-forward and recurrent pathways across higher-order areas of mouse visual cortex

**DOI:** 10.1101/2020.09.24.312140

**Authors:** Jennifer Y. Li, Charles A. Hass, Ian Matthews, Amy C. Kristl, Lindsey L. Glickfeld

## Abstract

Cortical visual processing transforms features of the external world into increasingly complex and specialized neuronal representations. These transformations arise in part through target-specific routing of information; however, within-area computations may also contribute to area-specific function. Here, we sought to determine whether higher-order visual cortical areas LM, AL, PM, and AM have specialized anatomical and physiological properties by using a combination of whole-cell recordings and optogenetic stimulation of V1 axons *in vitro*. We discovered area-specific differences in the strength of recruitment of interneurons through feed-forward and recurrent pathways, as well as differences in cell-intrinsic properties and interneuron densities. These differences were most striking when comparing across medial and lateral areas, suggesting that these areas have distinct profiles for net excitability and integration of V1 inputs. Thus, cortical areas are not defined simply by the information they receive, but also by area-specific circuit properties that enable specialized filtering of these inputs.

## Introduction

The neocortex is involved in a diverse array of computations that are fundamental to sensory and motor processing. It encompasses areas that represent basic features like the contrast of a visual stimulus or the frequency of a tone, and also areas directly linked to decision-making, working memory, and motor planning (Douglas and Martin, 2004; Gnadt and Andersen, 1988; Rushworth et al., 2011). Given the wide range of these specialized functions, the overall structure of the areas that perform them is remarkably similar (Douglas and Martin, 2004; Tremblay et al., 2016). Neocortical areas are characterized by a six-layer architecture broadly comprised of two neuronal cell types: excitatory, glutamatergic pyramidal neurons and inhibitory, GABAergic interneurons. Further consideration of genetically defined subpopulations of these cell types and their connectivity profiles reveals specific motifs of circuit wiring. For example, feed-forward inhibition involving co-activation of pyramidal cells and interneurons is thought to involve parvalbumin-expressing (PV) interneurons, whereas late-onset, feedback inhibition is mediated by somatostatin-expressing (SOM) interneurons (Jang et al., 2020; Kapfer et al., 2007; Li et al., 2015; Ma et al., 2010). These types of feed-forward and feedback connections are observed throughout cortex and work in conjunction to combine hierarchical inputs across areas and local processing within areas (D’Souza et al., 2016; Douglas et al., 1995; Gabernet et al., 2005; Yang et al., 2013). Understanding how the repetition of basic cortical motifs can give rise to specialized representations is crucial for insight into how the brain transforms neuronal activity across areas to generate higher-order computations.

On one hand, the consistent patterning of cortical structure could translate to a similar uniformity in local processing. Sensory cortical areas have been shown to have the capacity to remap across modalities to compensate the loss of one; this flexibility of representation could arise from a large degree of anatomical and functional similarity of structure across areas (Rauschecker, 1995). In this model, higher order computations could be generated from specific arrangement of fundamental cortical units that are set up to route specific inputs. This seems appropriate in the case of macaque primary visual cortex (V1), where segregated projections to thin stripe and thick stripe/inter-stripe regions in V2 enable the parallel transmission of color and orientation signals, respectively (Livingstone and Hubel, 1988; Livingstone and Hubel, 1984). Similarly, motion information is selectively routed to MT through a subset of V1 neurons that are more likely than chance to be direction-selective (Movshon and Newsome, 1996). Yet, additional variation in individual components of these canonical cortical motifs could create specialized microcircuits that further tune responses within an area. Response timing and sensitivity have been shown to vary across different sensory cortical areas, even after controlling for sensory modality and subcortical activity (Yang and Zador, 2012). These specializations could be related to differences in intrinsic excitability of neurons, recurrent connectivity, and relative recruitment of excitation and inhibition as a function of neocortical topography (Fletcher and Williams, 2019; Luna and Pettit, 2010; Wang, 2020). However, evidence of specialization of multiple features across specific, functionally defined areas remains relatively sparse (D’Souza et al., 2016; Fulcher et al., 2019; Glickfeld and Olsen, 2017).

The visual system is an excellent model for understanding the relative contribution of different mechanisms to neocortical specialization. As information is transferred hierarchically between primary and higher visual areas (HVAs), there is divergence of visual representations from basic features such as orientation and contrast to objects and self-generated motion (Hubel and Wiesel, 1962; Niell and Stryker, 2008; Tanaka, 1996; Ungerleider and Haxby, 1994; Zemel and Sejnowski, 1998). Thus, visual perception involves the coordinated evolution of specialized components of a stimulus across many areas of the brain. While many studies of hierarchical transformation have largely been conducted in primates, recent studies have demonstrated similar functional specialization in areas of mouse cortex (Andermann et al., 2011; Glickfeld and Olsen, 2017; Marshel et al., 2011; Murgas et al., 2020; Sit and Goard, 2020).

In the mouse, V1 projects to many HVAs immediately surrounding it, including areas LM (lateromedial), AL (anterolateral), PM (posteromedial), and AM (anteromedial; Wang and Burkhalter, 2007). Similar to primate cortex, V1 projections to HVAs are selectively routed and are more likely to target areas with matched function. Consequently, different areas can share similar circuit motifs but have divergent functional properties depending on the inputs they receive (Glickfeld et al., 2013; Han et al., 2018; Matsui and Ohki, 2013). However, this divergence alone is insufficient to account for all differences in stimulus selectivity across areas (Blot et al., 2020; Glickfeld and Olsen, 2017; Murgas et al., 2020). The organization of HVAs around V1 separates many of these areas by millimeters of cortical space. Therefore, divergence could also arise from relatively localized specialization of cellular and circuit properties that imbue different HVAs with distinct computational capabilities (Fletcher and Williams, 2019; Kim et al., 2017; Wang, 2020).

To investigate these properties of mouse HVAs, we performed whole-cell patch clamp recordings of pyramidal cells and PV- and SOM-expressing subpopulations of interneurons in retinotopically matched regions of LM, AL, PM, and AM. We made systematic measurements of intrinsic properties of these cells, as well as features of their feed-forward and local connections using paired recordings and optogenetic stimulation. Our data demonstrate significant variation in anatomical and physiological properties across cortical areas, specifically across areas separated on medial versus lateral sides of V1. Compared to lateral areas, in medial areas PV cells were less dense, pyramidal cells had higher input resistance, and relative feed-forward input onto PV and SOM interneurons was lower. These trends indicate that the primary difference may be in the overall excitability between these areas. However, local connectivity between pyramidal cells and interneurons was higher in medial HVAs than lateral HVAs, suggesting this difference in feed-forward excitability could be compensated for by more recurrent recruitment of interneurons in these areas. Altogether, our results indicate that higher order visual areas are not uniform in their cellular and circuit properties and this may contribute to differences in response properties across areas.

## Results

### Functional identification of mouse HVAs in coronal slices

Typically, the HVAs are functionally identified using their signature retinotopic organization obtained via *in vivo* imaging approaches (Garrett et al., 2014). However, in order to investigate synaptic, cellular, and circuit properties of HVAs, it is important to accurately identify each area in coronal sections which are commonly used for histology and *in vitro* physiology. To make an atlas of the mouse HVAs in coronal slices, we combined functional imaging and histology within mice to compare *in vivo*, retinotopically identified positions of HVAs to their relative location *ex vivo* in the coronal slice (**Figure 1A**). We first identified the HVAs *in vivo* using intrinsic autofluorescence imaging (**Figure 1B**, left) and used this retinotopic map to target these areas for injection with fluorescent dyes (**Figure 1B**, right) before visualizing these injection sites *ex vivo* in coronal sections of the brain (**Figure 1C**). Across mice (n = 3), we found anatomical markers in coronal sections that allowed us to reliably identify four HVAs: LM (lateromedial), AL (anterolateral), PM (posteromedial) and AM (anteromedial). LM is the most posterior of these HVAs, and appears in sections with the superior colliculus and the most posterior portion of the hippocampus (before clear segregation of the dentate gyrus; **Figure 1C**, top); AL and PM are anterior to LM, typically in the same sections, at the level of the medial geniculate nucleus of the thalamus (**Figure 1C**, middle); AM is the most anterior of these HVAs, appearing at the level of the lateral geniculate nucleus of the thalamus (**Figure 1C**, bottom). We compared these to the projection pattern of V1 axons in coronal slices that were labeled via injection of the excitatory opsin oChIEF fused to the fluorescent reporter tdTomato (tdT) into V1 of EMX1-Cre mice (a line that expresses Cre in a broad population of excitatory neurons across layers; Figure 2A) (Gorski et al., 2002). Thus, when visualizing axons from V1 to the HVAs for both anatomy and physiology, we could use these landmarks to better distinguish areas along anterior-posterior axis and ultimately study retinotopically matched regions of these HVAs (**Figure 1D**).

**Figure 1.**
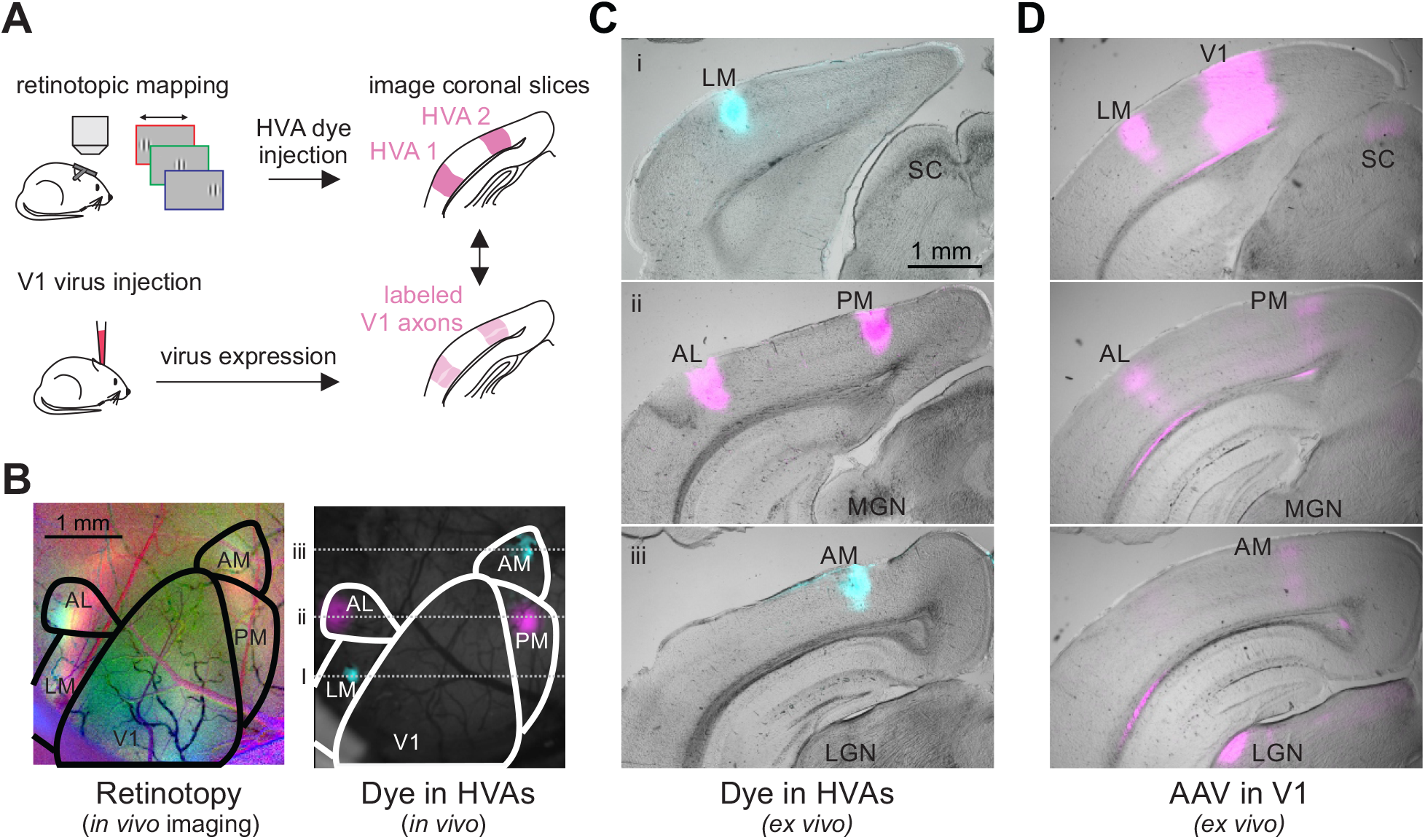
In vivo retinotopic mapping enables identification of HVAs in coronal slices. **A**. Schematic of procedure for identifying and labeling HVAs using both dye injections directly into the HVAs (top) and adeno-associated virus (AAV) mediated fluorophore expression in V1 axons (bottom). **B**. Left: Retinotopic map of left visual cortex with stimuli presented at 3 positions (azimuth: −10, red; +10, green; +30, blue). Right: Same field of view as on left, with blue and magenta dye injections in LM/AM and AL/PM, respectively. **C**. Coronal sections from the brain in **B** ordered from posterior (top) to anterior (bottom) with HVAs and other landmarks labeled (SC = superior colliculus; MGN = medial geniculate nucleus; LGN = lateral geniculate nucleus). Locations of coronal sections in anterior-posterior axis correspond to dotted lines in **B. D**. Coronal sections, at the same anterior-posterior locations as in **C**, from a different mouse with viral fluorophore expression in V1 neurons and their axons in the HVAs and other target regions (SC, LGN). Note the alignment of the V1 axon arborizations with the areas labeled via dye injection in **C**.

**Figure 2.**
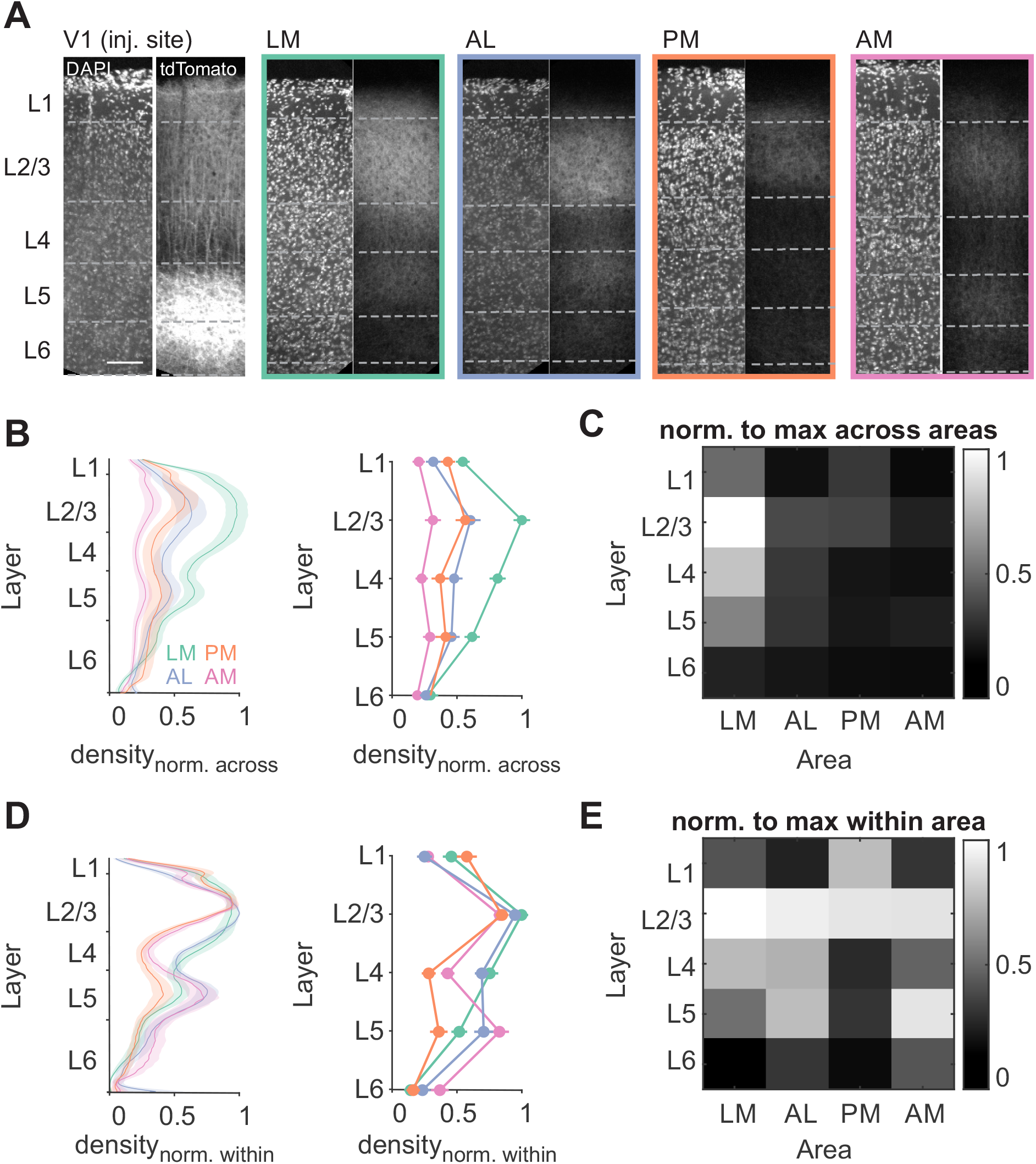
V1 axon densities are distinct across layers and HVAs. **A**. Confocal images of V1 and HVAs from an example EMX-Cre mouse injected with an AAV driving Cre-dependent oChIEF-tdTomato expression in V1 neurons. Dashed lines indicate layer boundaries as defined by DAPI staining. Scale bar = 100 µm. **B**. Fluorescence density by depth (left) or binned by layer (right), normalized to maximum fluorescence across all areas (typically in L2/3 of LM). Error bars are SEM across mice (n = 6 mice; LM-green, AL-orange, PM-blue, AM-pink). **C**. Heatmap summary of **B** (right). **D-E** Same as **B-C**, normalized to maximum density within each area.

One key determinant of the integration of V1 input in the HVAs is the density and laminar distribution of these projections. Using this approach to study retinotopically matched regions of HVAs, we found that V1 axons are densest in area LM and in layer 2/3, consistent with previous studies (two-way ANOVA, LM vs all other areas: p<0.01, AL vs AM: p<0.01; L2/3 vs all other layers: p<0.05; n = 8 mice; **Figure 2B-C**) (Wang, Sporns, & Burkhalter, 2012; Yang et al., 2013). However, in addition to these general trends, we also find significant differences in laminar distribution across areas. Inputs to layer 5 of AM and layer 1 of PM are denser than projections to these layers in other areas (**Figure 2D-E**; Layer 5: AM vs LM p<0.01, AM vs PM p<0.001; Layer 1: AL vs PM, p<0.01), and a higher fraction of inputs target layer 4 of lateral areas than medial areas (LM vs PM, p<0.001, LM vs AM p<0.05. AL vs PM p<0.001, AL vs AM p<0.05). Differences across areas were generally larger than individual variability within areas (**Figure 2 – figure supplement 1**). Thus, differences in the density and distribution of V1 axons across areas suggests heterogeneity in the impact of V1 on its targets. Moreover, the match between these data and the literature indicate that this viral-expression/landmark-based method consistently and accurately identifies the HVAs in coronal sections.

### Layer-specific differences in density of inhibitory interneurons across HVAs

Using this method to identify HVAs in the mouse, we next addressed whether different HVAs have distinct biological properties that could contribute to distinct stimulus preferences *in vivo*. Synaptic inhibition provided by local inhibitory interneurons is a critical factor in determining the time scale of integration in cortical neurons and their visual response properties (Adesnik, 2017; Adesnik et al., 2012; Priebe and Ferster, 2008; Reinhold et al., 2015; Wilson et al., 2018). Any differences in the relative proportions, recruitment, or connectivity of PV and SOM cells can influence integration and visual response properties in the HVAs (Kim et al., 2017). Thus, we first used an anatomical approach to measure the relative densities of PV and SOM interneurons in the HVAs.

Comparing each of these inhibitory subpopulations across areas, we found that inhibitory networks differ anatomically across medial and lateral areas. To label PV or SOM cells, we crossed PV-Cre or SOM-Cre mice with Ai14 reporter mice to drive tdT expression. In each mouse, we virally expressed Chronos-GFP in V1 and used the axonal arborizations to delineate retinotopically matched regions of the HVAs. These boundaries were then used to calculate the density of interneurons in each area (**Figure 3A**). This analysis reveals that PV cells are significantly more numerous in L4 and L5 of LM and AL, while SOM cells are more numerous in L2/3 and L6 of PM and AM (PV: one-way ANOVA with post hoc Tukey tests; L4 p<0.05, AL vs AM, p<0.05; L5 p<0.05; LM vs PM p<0.05, LM vs AM p<0.05; n=8 mice; SOM: one-way ANOVA with post hoc Tukey tests; L2/3 p<0.001, LM vs AM p<0.01, AL vs AM, p<0.01; L6 p<0.05, LM vs PM p<0.05, AL vs PM, p<0.05; n=5 mice; **Figure 3B**). Notably, the relative ratio of PV and SOM cells is more similar within layers than across areas (distance within layer, across areas: 0.56±0.31, distance within areas, across layers: 2.11±0.73, p<0.001 **Figure 3C, Figure 3 – figure supplement 1**), such that major trends hold across areas: relative enrichment of PV cells compared to SOM cells in L4, relative sparsity of PV and SOM cells in L2/3, and relative enrichment of PV and SOM cells in L5. However, within each layer, areas that share the same medial-lateral axis (i.e. LM/AL and PM/AM) are more similar than those that share the same anterior-posterior axis (i.e. LM/PM and AL/AM) or share neither axis (i.e. LM/AM and PM/AL) (**Figure 3D**; one-way ANOVA with post hoc Tukey tests; shared M-L vs shared A-P, p<0.01; shared M-L vs neither axis, p<0.01). Together, these differences in density of both PV and SOM cells across layers may shape excitability in an area- and layer-specific manner.

**Figure 3.**
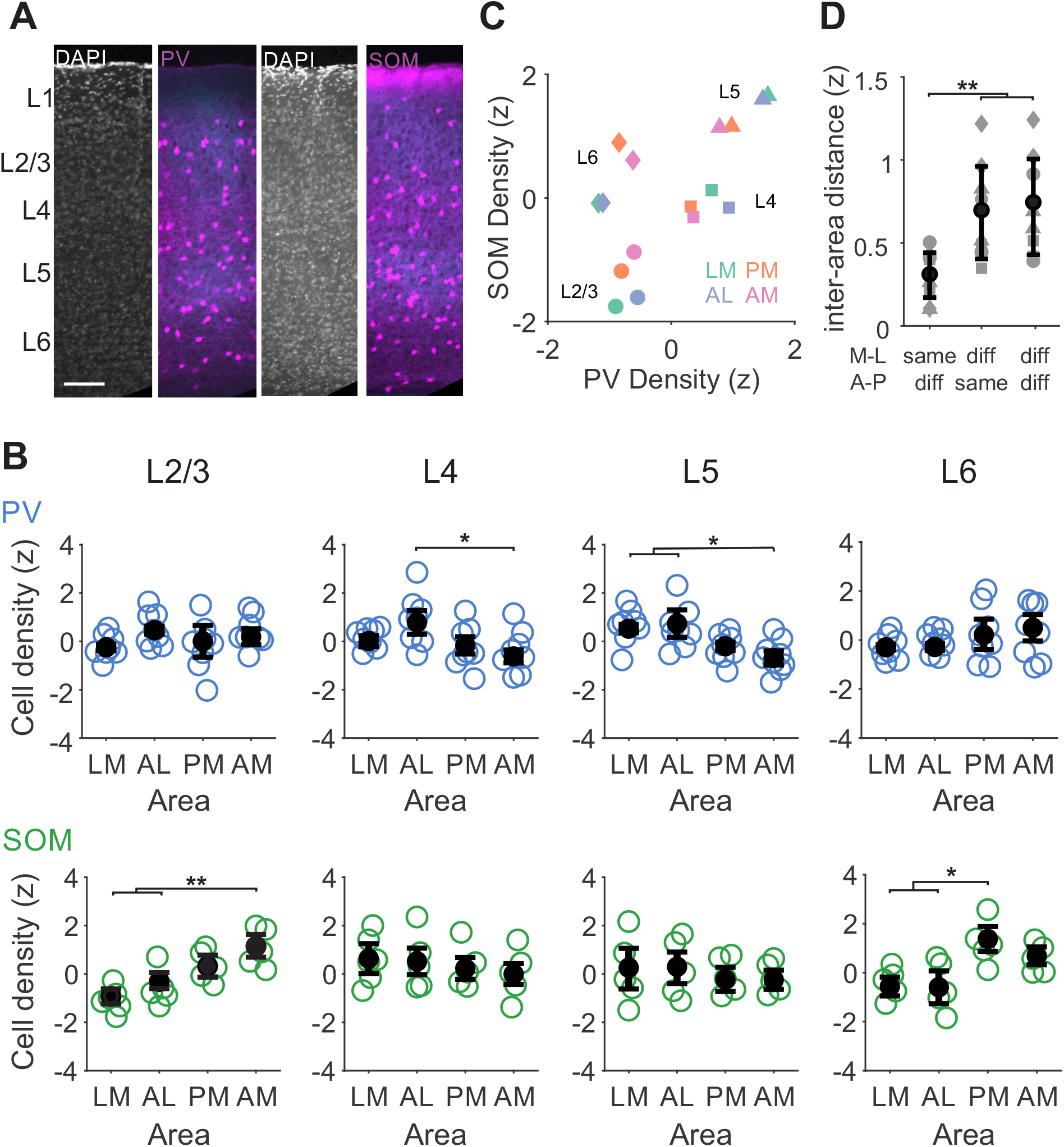
Different relative densities of PV and SOM interneurons across layers and HVAs. **A**. Confocal images from two example mice with either PV (left) or SOM (right) interneurons transgenically labeled with tdTomato (tdT; magenta) and V1 axons virally labeled with Chronos-GFP (blue). Layer boundaries are defined using a DAPI stain. **B**. Top, blue: Z-scored PV cell densities across HVAs for each layer. *, **, and *** denote p < 0.05, 0.01, and 0.001 (n = 8 mice). Bottom, green: Same as top for SOM cells (n = 5 mice). Open circles-individual mice; closed circles-average across mice. Error bars are SEM across mice. **C**. Scatter plot of z-scored PV and SOM densities. Each marker represents the average density across all mice of each cell type. Shape denotes layer, color indicates area. Note that distances between layers are shorter than distances between areas. **D**. Euclidian distance between points in **C** within the same layer, grouped by medial or lateral areas (LM-AL and PM-AM), anterior or posterior areas (LM-PM and AL-AM), or neither (LM-AM and AL-PM). Each point is a pairwise distance within layer (symbols corresponding to same layer shapes as **C**). Error bars are SD across layers.

### Intrinsic properties of pyramidal cells and interneurons are similar across HVAs

Activity within a region is also influenced by neurons’ intrinsic membrane properties, which determine how they integrate inputs and fire action potentials (Bean, 2007; Connors et al., 1982; Yamashita et al., 2013). These properties differ significantly between cell types, such as between pyramidal cells and interneurons, but also within cell type as a function of layer, cortical region, and projection target (Fletcher and Williams, 2019; Hu et al., 2014; Riedemann, 2019; Tremblay et al., 2016; Yamashita et al., 2013). Since these intrinsic factors ultimately determine synaptic integration, we next compared the intrinsic properties of both pyramidal cells and interneurons across HVAs.

We targeted pyramidal, PV and SOM cells within layer 2/3 of each HVA using a combination of the viral-expression based approach to identify HVAs (**Figure 1**) and mice with PV or SOM interneurons transgenically labeled with GFP to target these specific interneuron types. We performed whole-cell current clamp recordings while injecting current steps and measured membrane properties and spiking responses of neurons (**Figure 4A**). Unlabeled neurons with clear apical dendrites tend to have broad action potentials, strong spike frequency adaptation, and a small I_h_ sag due to the activation of a hyperpolarization-activated depolarizing current (**Figure 4B-C**; **Table 1**) consistent with these being excitatory pyramidal cells. In comparison, PV interneurons have narrow action potentials, marginal spike frequency adaptation, and less I_h_ sag, while SOM interneurons have slightly broader action potentials, moderate spike frequency adaptation, and a large I_h_ sag. Furthermore, SOM cells have characteristically high input resistance and a slow membrane time constant, whereas pyramidal cells and PV cells have lower input resistance as well as faster membrane time constants (**Figure 4D**; **Table 1**). These intrinsic properties result in significant differences in the input-output functions of each cell type as measured by spiking in response to depolarizing current injections—pyramidal cells’ broad action potentials and strong spike frequency adaptation led to relatively shallow F-I curves whereas PV cells’ F-I curves had steeper slopes related to narrower action potentials and less spike frequency adaptation, and SOM cells’ F-I curves demonstrated lower spike threshold associated with high input resistance (3-way ANOVA, main effect of I_inj_: p<0.001; main effect of cell type: p<0.05; interaction between current injection and cell type: p<0.05; **Figure 4E**). Altogether, our results are consistent with previous measurements of these cell types and demonstrate significant differences in characteristic features of neurons’ intrinsic properties for each of these three major cell types in the HVAs (Hu et al., 2014; Riedemann, 2019); 2-way ANOVA for spike frequency adaptation, half-width, I_h_, R_in_, and tau; main effect of cell type: p<0.05).

**Table 1.** Cell-intrinsic properties of Pyr, PV, and SOM cells across all HVAs. All data are mean ± SD.

|  | **Pyr** (n = 153) | **PV** (n = 66) | **SOM** (n = 56) |
| --- | --- | --- | --- |
| $R_{in}$ (MΩ) | 87.79 ± 26.31 | 102.07 ± 34.19 | 135.05 ± 35.49 |
| AP width (ms) | 0.83 ± 0.21 | 0.31 ± 0.08 | 0.49 ± 0.10 |
| AP adapt | 0.18 ± 0.05 | 0.81 ± 0.12 | 0.46 ± 0.12 |
| Membrane tau (ms) | 8.90 ± 2.85 | 7.98 ± 2.29 | 22.06 ± 8.37 |
| $I_h$ (μV/pA) | 4.43 ± 2.27 | 1.90 ± 0.92 | 8.05 ± 3.46 |

**Figure 4.**
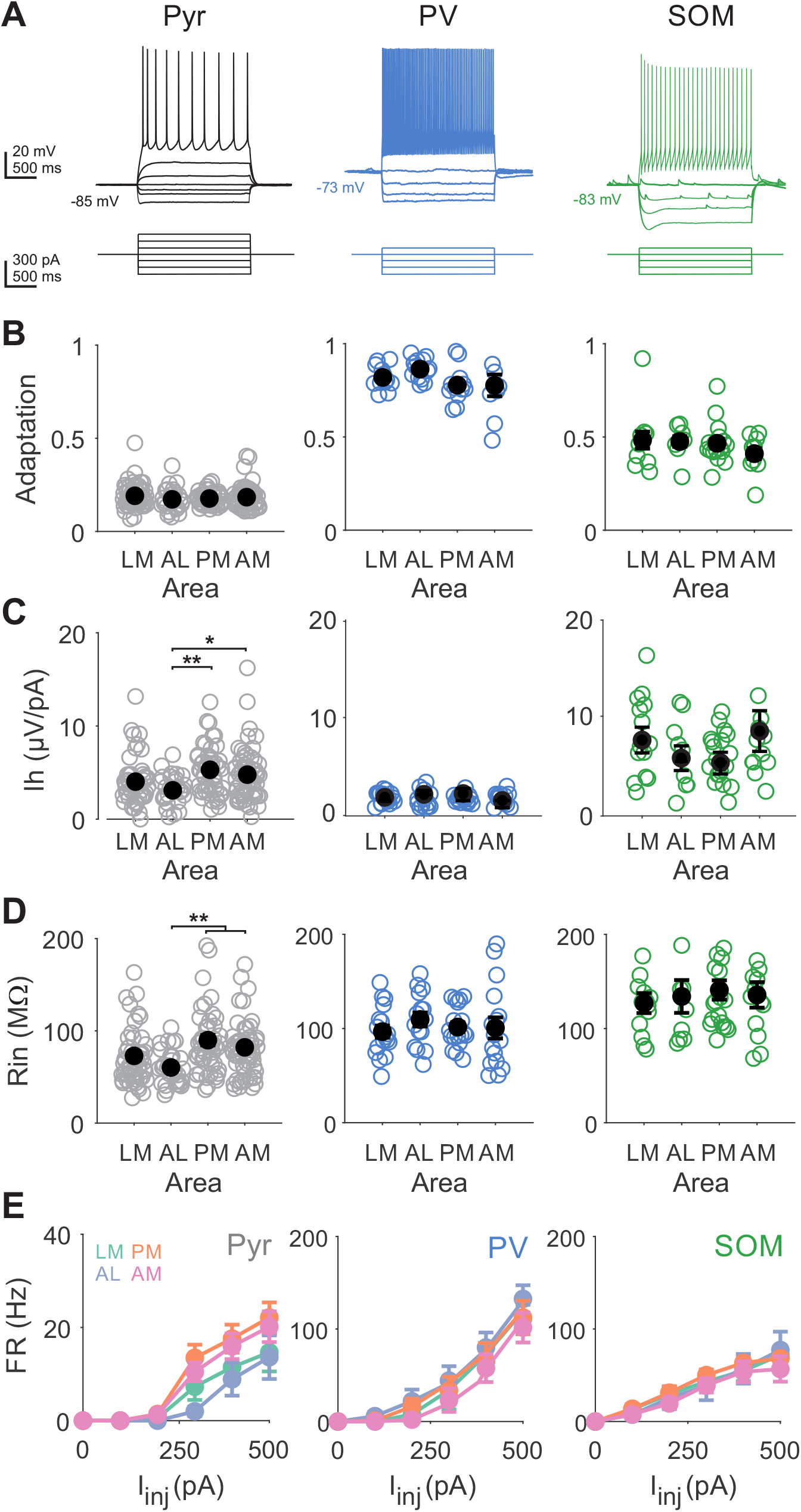
Intrinsic properties of pyramidal cells, but not interneurons, are distinct across areas. **A**. Example Pyr (black; left), PV (blue; center), and SOM (green; right) voltage traces (top) in response to current steps (bottom). All depolarizing steps until first spikes are shown. **B**. Spike rate adaptation (ISI_last_/ISI_first_) for pyramidal cells (LM = 55 cells, AL = 31, PM = 50, AM = 62), PV interneurons (LM = 17 cells, AL = 12, PM = 17, AM = 16), and SOM interneurons (LM = 15 cells, AL = 11, PM = 16, AM = 13) measured as slope of V-I curve around rest. Open circles-individual cells; closed circles-average across cells. Error bars are SEM across cells. *, **, and *** denote p < 0.05, 0.01, and 0.001. **C**. Same as **B**, for for voltage sag amplitude from activation of the hyperpolarization-activated current (I_h_). **D**. Same as **B**, for input resistance (R_in_). **E**. Firing rate (FR) vs current injection (I_inj_) for pyramidal cells (left), PV interneurons (center), and SOM interneurons (right) for each area. Error bars are SEM across cells.

While we find significant differences across cell types, most intrinsic properties are consistent across areas—spike frequency adaptation and membrane time constant are not significantly different across areas for pyramidal, PV or SOM cells, and PV and SOM cells have no significant difference in input resistance and I_h_ across HVAs (**Figure 4B-D**, center and right; main effect of area: p>0.05). The only differences across areas are the input resistance and I_h_ of pyramidal neurons in the medial versus lateral HVAs (**Figure 4B**, R_in_: AL vs PM p < 0.01, AL vs AM p <0.01; **Figure 4D**, I_h_: AL vs PM p<0.01, AL vs AM p<0.05). Although differences in cortical thickness across cortex have been shown to create gradients in input resistance (Fletcher and Williams, 2019), this difference was still present when between-area comparisons were restricted to measurements recorded within the same depths (LM vs PM p <0.05; LM vs AM p <0.05). Consistent with these differences in input resistance, there is also a significant difference in the input-output functions of pyramidal cells across areas (2-way ANOVA, main effect of I_inj_: p<0.001 main effect of area: p<0.01; **Figure 4E**), suggesting area-specific differences in excitability. Collectively, these data demonstrate that pyramidal cells across medial and lateral areas have differences in cell-intrinsic excitability, but most intrinsic passive and active properties of both excitatory and inhibitory neurons within layer 2/3 are relatively consistent between HVAs, suggesting similar filtering functions of V1 inputs.

### Area-specific differences in the excitation of interneurons and pyramidal cells by V1

Parallel excitatory input onto pyramidal cells and interneurons ties the strength and timing of local inhibition to that of feed-forward excitation in a process called feed-forward inhibition (Beierlein et al., 2003; Pouille and Scanziani, 2004; Tremblay et al., 2016). Typically, in primary sensory areas, feed-forward inputs drive PV cells more strongly than neighboring pyramidal cells such that PV cells provide robust, short-latency feed-forward inhibition (Gabernet et al., 2005). Notably, SOM cells generally receive weaker feed-forward excitation, and thus are not as robustly engaged in generating feed-forward inhibition (Hu and Agmon, 2016; Ma et al., 2010; Silberberg and Markram, 2007; Yavorska and Wehr, 2016). Thus, the relative strength of feed-forward V1 excitation onto different cell types is an important mechanism for regulating the balance and timing of excitation and inhibition within an area.

To directly compare the relative strength of V1 inputs across cell types and areas, we made simultaneous whole-cell voltage-clamp recordings from layer 2/3 pyramidal cells and interneurons while optogenetically stimulating V1 inputs (**Figure 5A-B**). Cells were voltage clamped at the reversal potential for inhibition to isolate excitatory post-synaptic currents (EPSCs). Stimulation of V1 axons sometimes evoked multiphasic EPSCs that likely reflect monosynaptic V1 excitation followed by recruitment of local recurrent circuits. To specifically isolate feed-forward V1 input, we used the first local minimum in the synaptic response (using the first minimum of the derivative of the current to find the steepest rise of the EPSC) to define a short window for finding the maximum EPSC response (Maier et al., 2011) (**Figure 5B**). This approach identified EPSCs with short latency and fast rise time in all three cell types, consistent with these EPSCs being due to monosynaptic V1 excitation (**Figure 5C-D**). EPSCs onto PV interneurons have significantly faster rise times and lower jitter than both pyramidal cells and SOM interneurons, but none of these measures of kinetics are significantly different across areas (rise time: two-way ANOVA with post hoc Tukey test; effect of area p>0.05, effect of cell type p<0.001; PV vs Pyr, p<0.001; PV vs SOM p<0.001, SOM vs Pyr p>0.05; jitter: two-way ANOVA with post hoc Tukey test; effect of area p>0.05, effect of cell type p<0.001; PV vs Pyr, p<0.001; PV vs SOM p<0.001; SOM vs Pyr, p<0.05).

**Figure 5.**
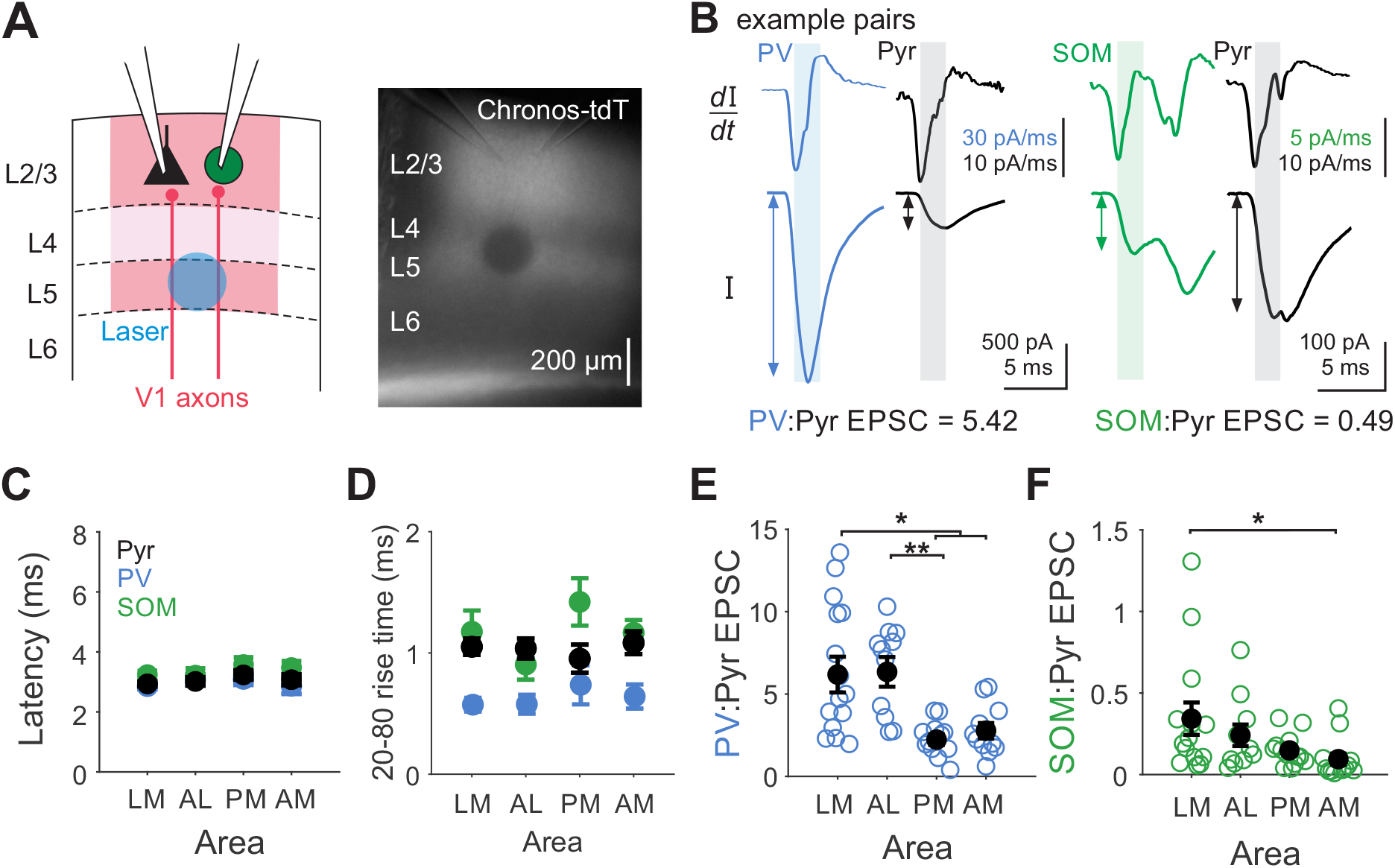
Different ratios of excitation onto pyramidal cells and interneurons across HVAs. **A**. Left: Schematic depicting setup for paired interneuron (IN) and Pyr recording with L5 optogenetic stimulation of V1 axons expressing Chronos. Right: Example image from slice with recording pipettes positioned in L2/3 and laser site photobleached after recording. **B**. Example EPSCs from V1 axon stimulation recorded in a PV and Pyr pair (PV blue, Pyr black; left) and a SOM and Pyr pair (SOM green, Pyr black; right). Top: First derivative of EPSCs (*dI/dt*). Short analysis window after derivative minimum (shaded region) is used to define early EPSC maximum search window. Bottom: Maximum EPSC amplitude in the early component of the response is measured for both IN and Pyr. **C**. Latency of EPSCs measured as time to 20% of maximum current. No significant effect of area or cell type (p>0.05). Error bars are SEM across cells. **D**. 20-80% rise time of EPSCs by cell type. No significant effect of area, significant effect of cell type (PV vs Pyr, p<0.001; PV vs SOM, p<0.001; two-way ANOVA with post-hoc Tukey test). **E**. Ratio of excitation onto each IN and pyramidal cell (IN:Pyr ratio) for pairs divided by HVA. Open circles-individual pairs; closed circles-average across pairs. Error is SEM across pairs. Left: PV:Pyr; LM = 15 pairs, AL = 11, PM = 12, AM = 11; right: SOM:Pyr; LM = 18 pairs, AL = 12, PM = 13, AM = 13. *, **, and *** denote p < 0.05, 0.01, and 0.001.

Similar to what has been observed at thalamocortical inputs onto V1 neurons, V1 inputs onto PV cells are significantly larger than onto paired pyramidal cells across all areas (p<0.001; paired t-test; n=49 pairs), while V1 inputs onto SOM cells are significantly weaker (p<0.001; paired t-test; n=56 pairs). In order to compare the relative excitation of cell types across areas, we normalized the EPSC onto each interneuron by its paired pyramidal cell to generate an IN:Pyr ratio (**Figure 5E**). Lateral areas tend to have larger IN:Pyr ratios than medial areas for both PV (p<0.05; one-way ANOVA; **Figure 5E**, left; LM vs PM or AM, p<0.05; AL vs PM p<0.05) and SOM interneurons (LM vs AM p<0.05; one-way ANOVA with post hoc Tukey test; **Figure 5E**, right). Importantly, this effect is not dependent on the strength of activation: when varying the laser strength within recordings of a pair, we find the ratio to be independent of the laser intensity for both PV (p>0.05; n=10 pairs; one-way ANOVA) and SOM cells (p>0.05; n=6 pairs; one-way ANOVA). This suggests that inhibitory interneurons are more strongly activated by the same set of V1 inputs in lateral areas as compared to medial areas.

### Late-onset, local recruitment of excitation differs across HVAs

When stimulating V1 axons, we also observed late-onset excitation that could arise from recruitment of recurrent circuits within the HVAs (**Figure 6A**) (Seeman et al., 2018). To quantify the relative recruitment of feed-forward and local excitation, we measured the time to half-max for the cumulative excitatory charge (**Figure 6B-C**). Shorter times reflect fast decaying EPSCs in which the majority of the current is monosynaptic, while longer times likely reflect the recruitment of polysynaptic inputs. EPSCs recorded in pyramidal cells have similar times to half-max across areas (**Figure 6D**, left), suggesting a similar proportion of feed-forward and local excitation. In comparison, EPSCs recorded in PV cells and SOM cells have different proportions of feed-forward and local input in medial compared to lateral areas. EPSCs onto SOM cells in PM and AM have significantly longer times to half-max than those in LM and AL (**Figure 6D**, right; PM vs LM: p < 0.05; PM vs AL: p < 0.01; AM vs AL: p < 0.05). A similar, though non-significant, trend is seen in PV cells (**Figure 6D**, center). Importantly, this difference in the time course cannot be explained by differences in EPSC kinetics, since EPSC rise times are not significantly different across areas (**Figure 5C-D**). This difference also cannot be attributed to differences in effective stimulation intensity of V1 axons, as EPSC amplitudes onto pyramidal cells were consistent across areas (**Figure 6 – figure supplement 1**). Indeed, inspection of the average EPSC traces sorted by area reveals that our measure reflects differences in the recruitment of secondary, longer latency EPSCs (**Figure 6E**). While the average time courses of EPSCs onto pyramidal cells are similar across medial and lateral areas, the average responses recorded in PV and SOM cells reveal later-onset excitation selectively in medial HVAs in both cell types. Thus, stimulation of V1 inputs to the HVAs differentially recruits local networks in medial versus lateral areas.

**Figure 6.**
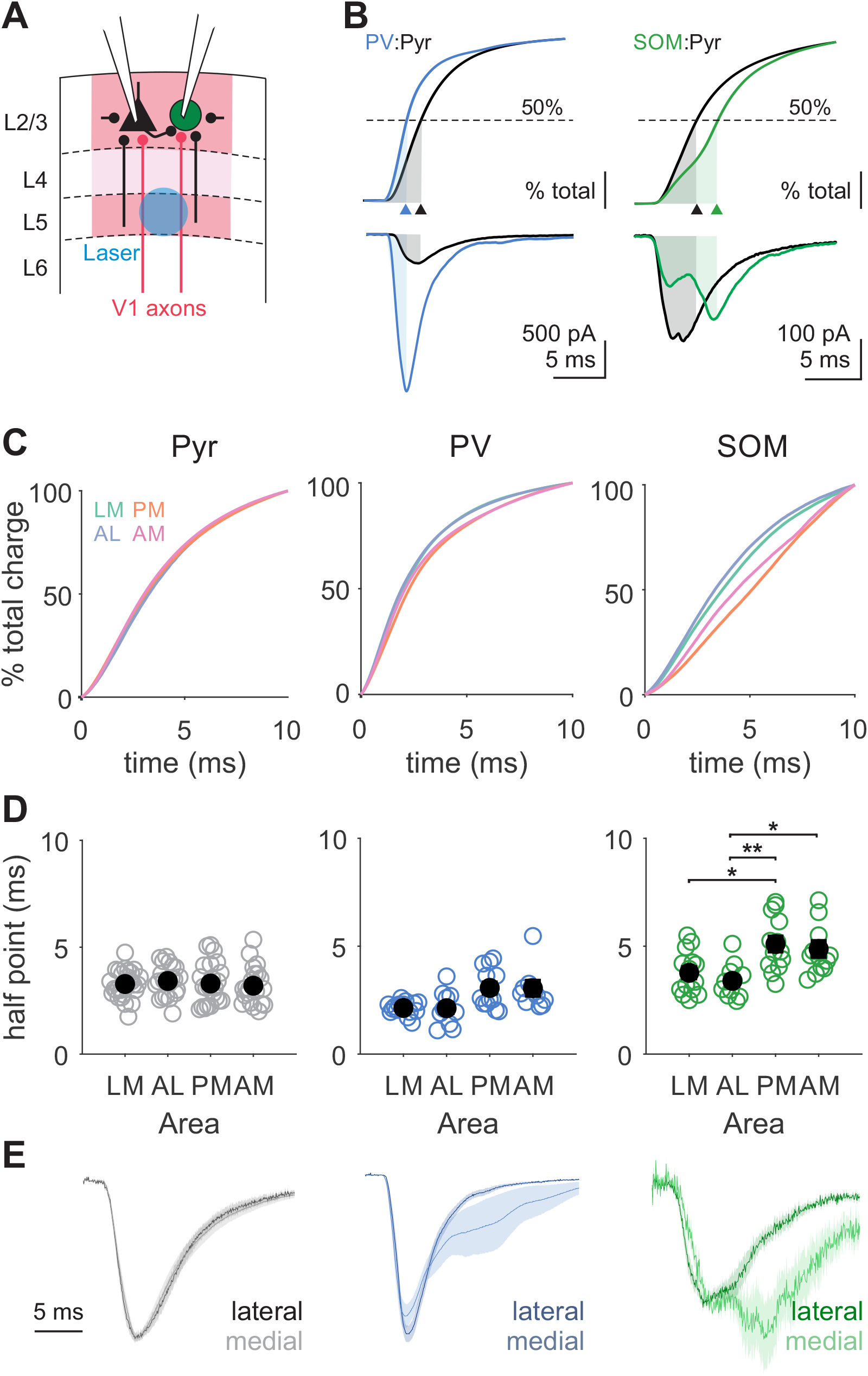
Interneurons in medial areas receive more late-onset, polysynaptic excitation. **A**. Schematic depicting setup for paired recordings in HVAs with local excitatory inputs within the HVA. Local inputs from neighboring pyramidal cells come from both within L2/3 and across layers (black lines). (B) Top: Cumulative charge for a PV (blue) and Pyr (black) pair (left) and a SOM (green) and Pyr (black) pair (right) with time to 50% charge labeled (shaded region, arrowhead). Bottom: EPSCs used for analysis in top. Same example cells as in **Figure 5B. C**. Grand average of cumulative charges for Pyr (left), PV (center), and SOM (right) cell types. **D**. Summary of time to 50% charge for each cell type. Open circles-individual cells; closed circles-average across cells. Error is SEM across cells. Pyr: LM = 33 cells, AL = 23, PM = 25, AM = 24; PV: LM = 15 cells, AL = 11, PM = 12, AM = 11; SOM: LM = 18 cells, AL = 12, PM = 13, AM = 13. *, **, and *** denote p < 0.05, 0.01, and 0.001. **E**. Normalized EPSCs averaged across neurons, grouped by lateral (dark; LM and AL) and medial (light; PM and AM) areas. Shaded error is SEM across cells.

### Different connectivity probabilities between L2/3 pyramidal cells and interneurons across HVAs

There are two possible mechanisms that might explain the stronger local input onto interneurons in medial HVAs. One possibility is that stimulation of V1 inputs to medial areas drives stronger activation of local circuits. This is consistent with the comparatively weaker feed-forward excitation onto interneurons which could enable increased excitability in medial areas. However, if overall activity was higher in medial areas, we would also expect to see a similar increase in local inputs onto pyramidal cells. Since this is not the case, it instead suggests that there might be a difference in the strength or probability of local excitation onto interneurons within the HVAs.

To test this hypothesis, we made paired recordings from neighboring layer 2/3 pyramidal cells and interneurons within ∼40 µm of each other (**Figure 7A**). The amplitude and short-term dynamics of these connections are consistent with what was expected for these cell types, but do not significantly vary by area. We find that pyramidal cells receive significantly stronger inhibition from PV cells than SOM cells but provide equal excitation to both cell types (one-way ANOVA PV→Pyr vs SOM→Pyr: p<0.05; Pyr→PV vs Pyr→SOM: p > 0.05; **Table 2**). However, when dividing by areas we find no significant differences between areas in the amplitude of the unitary EPSCs or IPSCs recorded in interneurons or pyramidal cells, respectively, for either PV+Pyr or SOM+Pyr pairs (one-way ANOVA, all comparisons: p > 0.05; **Figure 7B-C**). Similarly, while short-term plasticity of unitary connections depends on interneuron type (one-way ANOVA, PV vs SOM IN→Pyr-P2/P1: p < 0.05; P10/P1: p < 0.05; **Table 2**), there are no differences in the paired-pulse ratios within cell types between areas (one-way ANOVA, P2/P1 and P10/P1: p>0.05 for all area comparisons). Thus, the strength and short-term plasticity of connectivity between pyramidal cells and interneurons is consistent across different HVAs and comparable to that seen in primary sensory areas (Avermann et al., 2012; Beierlein et al., 2003; Kapfer et al., 2007; Seeman et al., 2018; Tremblay et al., 2016).

**Table 2.** Amplitude and paired-pulse ratio of Pyr and IN connections across all HVAs. All data are mean ± SEM.

|  | **Pyr → PV** | **Pyr → SOM** | **PV → Pyr** | **SOM → Pyr** |
| --- | --- | --- | --- | --- |
| Connected/Total | 34/82 = 0.41 | 28/60 = 0.47 | 55/82 = 0.67 | 43/60 = 0.71 |
| P1 amplitude (pA) | -20.32 ± 2.74 | -18.41 ± 4.56 | 21.65 ± 1.79 | 15.05 ± 1.53 |
| P2/P1 | 0.95 ± 0.08 | 0.96 ± 0.09 | 0.87 ± 0.03 | 0.90 ± 0.03 |
| P10/P1 | 0.86 ± 0.05 | 1.69 ± 0.18 | 0.74 ± 0.03 | 0.93 ± 0.05 |

**Figure 7.**
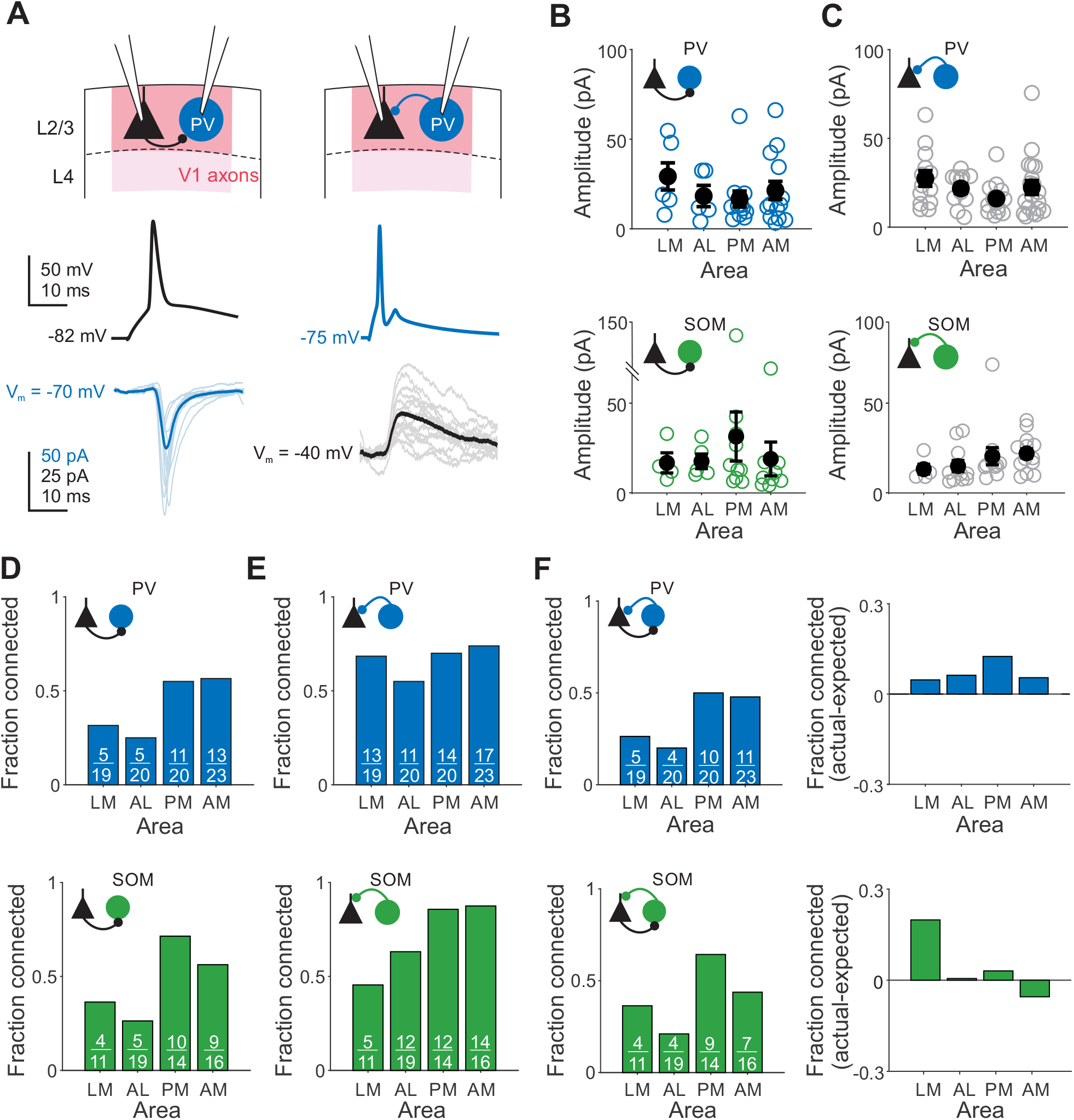
Distinct local connectivity between pyramidal cells and interneurons across HVAs. **A**. Example PV (blue) and Pyr (black) pair with both Pyr→IN connection (left) and IN→Pyr connection (right). Thin lines are individual trials, thick lines are mean. **B**. Amplitude of monosynaptic connections for Pyr→PV inputs (top) or Pyr→SOM inputs (bottom). Open circles-individual pairs; closed circles-average across pairs. Error bars are SEM across pairs. **C**. Same as **B**, for IN→Pyr connections. **D**. Fraction of connected Pyr→PV (top) and Pyr→SOM (bottom) pairs. **E**. Same as **D**, for IN→Pyr pairs. **F**. Left: same as **D**, for bidirectionally connected IN↔Pyr pairs. Right: Difference between actual fraction of bidirectionally connected pairs and expected fraction [P(Pyr→IN)*P(IN→Pyr)]. *, **, and *** denote p < 0.05, 0.01, and 0.001.

Although the amplitude and plasticity of connections do not vary as a function of area, we do find significant differences in the probability of these connections across areas. Both PV and SOM cells in medial areas are more likely than those in lateral areas to receive input from a neighboring pyramidal cell (Chi-squared test; Pyr→PV: p < 0.05; Pyr→SOM: p < 0.05**; Figure 7D**). SOM interneurons in medial areas are also more likely to inhibit neighboring pyramidal cells (Chi-squared test; SOM -> Pyr: p<0.05 **Figure 7E**), and both interneuron types are more likely to be bidirectionally coupled in medial areas (Chi-squared test; Pyr↔PV: p < 0.05; Pyr↔SOM: p < 0.05 **Figure 7F**, left). The probability of bidirectional connectivity is above chance for both PV and SOM cells: interneurons that receive excitatory input from a neighboring pyramidal cell are more likely to inhibit that pyramidal cell across all areas for PV cells, and exclusively but strongly in LM for SOM cells (**Figure 7F**, right). Altogether, these results reveal that different HVAs have distinct probabilities of local connectivity between interneurons and pyramidal cells within layer 2/3, with significantly higher probabilities in medial compared to lateral areas.

## Discussion

The neocortex is a highly conserved anatomical structure that performs a wide variety of computational tasks. Evidence suggests that specialization of function could arise through targeted routing of inputs to specific cortical areas (Churchland and Lisberger, 2005; Glickfeld et al., 2013; Livingstone and Hubel, 1988; Movshon and Newsome, 1996). However, some features of cellular and circuit properties have been shown to vary as a function of neocortical topography, such that areas have demonstrable differences along spatial axes (Fletcher and Williams, 2019; Kim et al., 2017; Wang, 2020; Yang et al., 2013). Here, we used a comparative approach to determine whether these differences could map onto a subset of visual areas that have divergent stimulus preferences (LM, AL, PM, AM) (Andermann et al., 2011; Glickfeld and Olsen, 2017). Utilizing a combination of anatomical and physiological techniques we found significant variation, primarily across medial versus lateral areas. The trends were indicative of a net difference in excitability across areas–in medial areas, PV interneurons were sparser (**Figure 3**), pyramidal cells had higher input resistance (**Figure 4**), and relative feed-forward input onto PV and SOM interneurons was lower (**Figure 5**). However, these differences may be compensated by complementary increases in local excitation of interneurons within the HVAs (**Figures 6-7**). Thus, in this study we have demonstrated that cortical microcircuits share motifs but are not mirror images of each other (**Figure 8**). Moreover, variation of these motifs could shape how inputs are filtered in an area and ultimately support specialization of cortical function.

**Figure 8.**
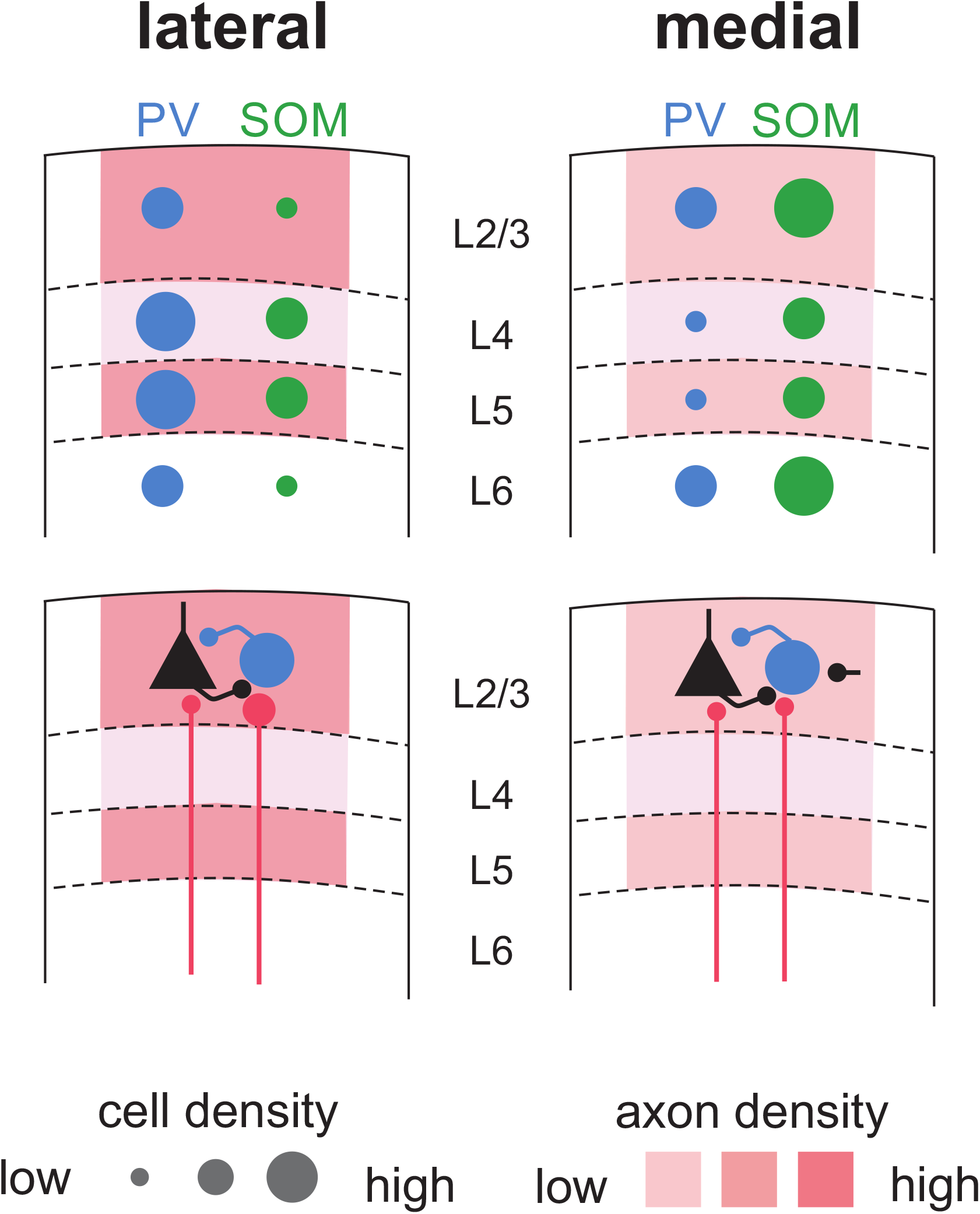
Lateral and medial HVAs have distinct anatomical and physiological profiles. Top: Schematic showing relatively higher density of axons, higher PV interneuron density in L4 and L5, and lower SOM interneuron density in L2/3 and L6 in lateral (left) compared to medial (right) areas. Bottom: Schematic showing relatively larger IN:Pyr EPSC ratio for feed-forward input (red inputs) but lower probability of local excitatory input (black inputs) in lateral (left) compared to medial (right) areas.

This spatial divide of cellular and circuit properties across medial and lateral areas is reminiscent of specialization of dorsal/ventral streams of visual processing in primates (Mishkin et al., 1983; Ungerleider and Haxby, 1994). Anatomical clustering of HVAs by the weights of inputs to each area is strongly correlated with the relative location of the areas in cortex, indicative of separate streams of processing (Wang et al., 2012). Our findings are consistent with studies separating areas PM and AM from LM, but depart from previous studies that distinguish AL and LM as parts of the dorsal and ventral streams, respectively (Wang et al., 2012, 2011; Wang and Burkhalter, 2013). Instead, we find properties of AL to be more similar to LM than PM or AM such that the segregation of function is biased to process stimuli more similarly within medial and lateral areas than across them. Notably, in the visual system, medial and lateral HVAs are separated by larger distances than those along the anterior-posterior axis. If these differences arise in part due to anatomical gradients (Fletcher and Williams, 2019; Kim et al., 2017; Wang, 2020), then larger medial-lateral distances might explain some of the bias that translates to segregated functional streams. Regardless of origin, in this study we have identified significant distinctions in intrinsic properties, interneuron density, and strength and distribution of feed-forward input across HVAs that could contribute to specialization of responses within an area.

Significant differences in these cell-intrinsic, synaptic, and circuit features could converge to generate broader differences in how inputs are transformed through the canonical circuit motifs present across neocortex. Feed-forward inhibition is a circuit motif that involves regulation of pyramidal cell activity with interneuron activity yoked through shared input (Douglas and Martin, 2004; Jang et al., 2020). Consistent with studies using other techniques to measure strength of feed-forward inputs in cortex, we found larger relative V1 input to PV cells than neighboring pyramidal cells (PV:Pyr > 1) and smaller input onto SOM interneurons than pyramidal cells (SOM:Pyr EPSC < 1) (D’Souza et al., 2016; Hu and Agmon, 2016; Yang et al., 2013). We also found an area-specific bias in these ratios, where both classes of interneurons in medial areas receive comparatively less excitation than neighboring pyramidal cells. This finding is different from previous studies measuring V1 inputs in HVAs using subcellular mapping, which found a larger PV:Pyr EPSC ratio in PM than LM (D’Souza et al., 2016; Yang et al., 2013). Here we evoked transmitter release in layer 2/3 via action potentials generated with axon stimulation at a single site in layer 5, whereas subcellular mapping involves localized, direct depolarization of axon terminals in tetrodotoxin (TTX) and 4-AP to block action potential generation and optimize terminal depolarization. These differences in stimulation could be a source of variability—whereas vesicle release associated with action potential generation is more physiological, it can also trigger recurrent inputs (**Figure 6**). While we have made an effort to isolate the early EPSC component and found that this component has similar kinetics to EPSCs recorded in TTX and 4AP (**Figure 5**), it is possible that some of the early input we measured is not solely from V1 axons. However, the stronger recurrent excitation of interneurons in medial areas should, if anything, bias our feed-forward measures to also be stronger in medial areas.

Excess excitation within a cortical circuit is detrimental not only from a metabolic perspective, but also detrimental for network stability (Isaacson and Scanziani, 2011; Ozeki et al., 2009; Sanzeni et al., 2020). Our results suggest that the weaker feed-forward inhibitory pathway in medial areas may be balanced in part by stronger feedback inhibitory pathways. SOM interneurons are implicated in this processing pathway (Adesnik, 2017; Kato et al., 2017; Priebe and Ferster, 2008), and were significantly denser in both L2/3 and L6 of medial areas (**Figure 3B**, bottom). Furthermore, we observed that interneurons in medial areas are more likely to receive late-onset excitation, likely arising from excitatory inputs from neighboring pyramidal cells that are driven to spike threshold by V1 inputs (**Figure 6**). This difference was most striking in SOM interneurons, but was also present in PV interneurons, and absent in pyramidal cells. This could be related to cell-type differences in recurrent connectivity, as SOM interneurons have been shown to be densely embedded in local networks (Fino and Yuste, 2011; Kapfer et al., 2007; Silberberg and Markram, 2007; Yavorska and Wehr, 2016), whereas pyramidal cells have comparatively sparse and specific recurrent connections (Cossell et al., 2015). This difference in late-onset charge appeared to arise from increased probability of recurrent excitatory inputs rather than larger amplitude of connections (**Figure 7B**). Together, our findings indicate that medial and lateral areas may be specialized in circuit features that converge to differentiate recruitment of feed-forward and feedback inhibitory pathways.

How could differences in cell-intrinsic and circuit properties across areas ultimately lead to the differences in stimulus specificity observed across these areas? Given what is known about the role of inhibition in stimulus tuning as well as relative contributions of feed-forward and recurrent circuitry, we can make several predictions on the net effect of the differences we have identified (Cardin, 2018; Douglas et al., 1995; Isaacson and Scanziani, 2011; Lien and Scanziani, 2013; Liu et al., 2011; Priebe and Ferster, 2008). Stronger feed-forward inhibition, primarily thought to be mediated by PV interneurons, improves representation of the timing of sensory input by narrowing the integration time window (Pouille and Scanziani, 2004). The effect of this is to reduce efficacy of asynchronous inputs that do not drive postsynaptic neurons to spike in the time preceding PV-mediated inhibition. Thus, stronger recruitment of feed-forward inhibition in lateral areas could improve encoding of higher frequency temporal features than medial areas. Indeed, neurons in AL have been shown to have higher temporal frequency preferences than neurons in PM (Andermann et al., 2011), and the stronger excitation of PV cells in AL could amplify the anatomical divergence in temporal frequency preferences of inputs to HVAs (Andermann et al., 2011; Glickfeld et al., 2013).

More generally, differences in recruitment of local inhibition may serve to alter excitability across areas and modulate response thresholds and sensitivity (Priebe and Ferster, 2008; Zhu et al., 2015). While no significant differences in contrast response functions have been observed across areas (Murgas et al., 2020), contrast normalization is thought to engage both feed-forward and feedback inhibition, and thus the opposing effects may result in little net effect on contrast responses (Carandini and Heeger, 2012; Nienborg et al., 2013). Instead, surround suppression thought to be generated specifically through feedback recruitment of SOM cells (Adesnik, 2017; Adesnik et al., 2012). Thus, given the stronger recruitment of SOM cells in medial areas, it is surprising that neurons in PM actually have larger receptive fields and less surround suppression than neurons in lateral areas (Murgas et al., 2020). However, spatial integration also relies on recurrent connections between pyramidal cells (Cossell et al., 2015; Douglas et al., 1995), as well as connections between pyramidal and SOM cells separated by hundreds of microns (Adesnik et al., 2012; Avermann et al., 2012; Fino and Yuste, 2011). Here, we have only tested connectivity of immediately neighboring pyramidal cells and interneurons, which we have shown to exhibit increased connection probability in medial areas. Further study of the probabilities for both of these types of connections over larger scales will be important to understand how connectivity patterns might contribute to spatial integration within an area.

In this study, we have generated evidence for specialization of cell-intrinsic, synaptic, and circuit features across multiple areas of the neocortex. By performing systematic measurements of various anatomical and physiological features across a subset of mouse higher visual areas, we have identified a medio-lateral division of local processing across higher-order visual areas that is similar to dorsal and ventral streams in other mammals. These findings are relevant not only for understanding how visual information is transformed across areas, but also highlight a set of potential mechanisms for larger-scale specialization of cortical computations across the brain.

## Methods

### Animals

All procedures conformed to standards set forth by the National Institutes of Health Guide for the Care and Use of Laboratory Animals, and were approved by the Duke University’s Animal Care and Use Committee. 143 mice (81 female) were used in this study. The following driver lines were used to express Cre-dependent optogenetic or fluorescent proteins: EMX-1-IRES-Cre (Jackson Labs #005628, n=63), SST-IRES-Cre (Jackson Labs #013044, n=40), PV-Cre (Jackson Labs #008069, n=40). To fluorescently label genetically identified interneurons, we crossed Cre-driver lines with Ai14 (Jackson Labs #007914) or Ai3 (Jackson Labs #007903) reporter mice. All transgenic mice were heterozygous and bred on a C57/B6J (Jackson Labs #000664) background. Mice were 28-40 days old at the time of injection and 32-60 days old at the time of sacrifice for histology or recording.

### Viral expression of exogenous proteins in V1 neurons

We expressed the light-gated cation channel oChIEF (Lin et al., 2009) or Chronos (Klapoetke et al., 2014) using recombinant adeno-associated viruses (rAAVs). Chronos was manufactured at the University of Pennsylvania Vector Core (AAV2/8.Syn.Chronos.tdTomato, Addgene 62726, 1.6×10^13^ GC/ml). oChIEF was manufactured by Virovek (AAV2/1.CAG.flex.oChIEF.tdTomato, Addgene 30541, 2.2×10^13^ GC/ml). For anatomical experiments, we expressed fluorophores using AAVs (AAV1.CB7.CI.eGFP.WPRE.rBG, University of Pennsylvania Vector Core CS0326, 2.04×10^13^ GC/ml; AAV8.Syn.Chronos.tdTomato, Addgene 62726, 1.6×10^13^ GC/ml; AAV9.CAG. flex.tdTomato.WPRE.bG, University of Pennsylvania Vector Core CS0476, 1.83×10^13^ GC/ml) in V1 to label projections to the HVAs to measure expression density and to facilitate area identification.

Viruses were pressure injected into the brain via a burr-hole. Briefly, isoflurane anesthetized mice were positioned in a stereotax (Kopf Instruments) and a small hole was drilled though the skull −2.6 mm lateral from lambda and directly anterior to the lambdoid suture (targeting the posterior and medial aspect of the primary visual cortex, V1). A glass micropipette containing the virus was mounted on a Hamilton syringe, lowered 350 – 500 µm into the brain, and 100 nL of virus or was pressure injected using an UltraMicroPump (World Precisions Instruments). We waited a minimum of 2 weeks for opsin expression.

### In vivo mapping for functional identification of HVAs in coronal sections

To find the correspondence between areas identified *in vivo* and in coronal section, mice were implanted with a headpost and a 5 mm cranial window (Goldey et al., 2014). After recovery from surgery, retinotopic maps were generated from intrinsic autofluorescence. The brain was illuminated with blue light 473 nm LED (Thorlabs), and emitted light was measured through a green and red filter (500 nm longpass). Images were collected using a CCD camera (Rolera EMC-2, Qimaging) at 2 Hz through a 5x air immersion objective (0.14 numerical aperture (NA), Mitutoyo), using Micromanager acquisition software (NIH). Visual stimuli were presented on a 144-Hz (Asus) LCD monitor, calibrated with an i1 Display Pro (X-rite). The monitor was positioned 21 cm from the contralateral eye. Visual stimuli were controlled with MWorks (http://mworks-project.org). Circular gabor patches (30° diameter) containing static sine-wave gratings alternated with periods of uniform mean luminance (60 cd/m^2^). Images were analyzed in ImageJ (NIH) to measure changes in fluorescence (dF/F; with F being the average of all frames) to identify primary visual cortex (V1) and the higher visual areas. Vascular landmarks were used to identify targeted sites for injection of Alexa-conjugated dextrans (488 or 594; 10,000 KD; Life Technologies). To inject the dye, the cranial window was transiently removed, and the same protocol used to inject virus was used to inject 50 nL dye (5% in water).

### Histology and imaging

Three weeks following the viral injection (or the day after for dye injections), mice were transcardially perfused with 4% paraformaldehyde. Whole brains were removed and post-fixed <24hrs in 4% paraformaldehyde. The paraformaldehyde was washed out of the tissue with 3×1hr rinses with PBS buffer containing (in mM): 137 NaCl, 2.7 KCl, 10 Na_2_HPO_4_, 1.8 KH_2_PO_4_. Fixed tissue was then soaked in a 30% sucrose solution until the tissue sunk. Brains were then sectioned in coronally (70 µm) using a freezing microtome and mounted onto microscope slides using a DAPI mounting medium (DAPI Fluoromount G, Southern Biotech).

Coronal brain slices were first imaged at 2x magnification using an epifluoresence microscope to determine the locations of the HVAs using a combination of the axonal arborizations of the V1 projections and anatomical landmarks. Next, confocal z-stacks (10x objective, 0,3 NA, Zeiss Axiovert 200M) were collected from the four consecutive slices with the densest V1 projections in each HVA. For measurements of projection density, the PMT gain was optimized for area LM (since this area generally has the strongest expression) and kept constant for all other areas to remove any variability due to differences in image acquisition.

### In vitro slice preparation and recordings

Mice were anesthetized with isoflurane, the brain was removed and then transferred to oxygenated (95% O_2_ and 5% CO_2_), ice-cold (0 – 4° C), artificial cerebrospinal fluid (ACSF, in mM: NaCl 126, KCl 2.5, NaHCO_3_ 26, NaH_2_PO_4_ 1.25, glucose 20, CaCl_2_ 2, MgCl_2_ 1.3). Coronal brain slices (350 µm) were prepared using a vibrating microtome (VT1200S, Leica) and transferred to a holding solution (at 34° C) for 12 minutes, and then transferred to storage solution for 30 min before being brought to room temperature. The holding solution contained (in mM): 92 NaCl, 2.5 KCl, 1.25 NaH_2_PO_4_, 30 NaHCO_3_, 20 HEPES, 25 glucose, 2 thiourea, 5 Na-ascorbate, 3 Na-pyruvate, 2 CaCl_2_, 2 MgSO_4_. The storage solution contained (in mM): 93 NMDG, 2.5 KCl, 1.2 NaH_2_PO_4_, 30 NaHCO_3_, 20 HEPES, 25 glucose, 2 thiourea, 5 Na-ascorbate, 3 Na-pyruvate, 0.5 CaCl_2_, 10 MgSO_4_. Intracellular recordings of were obtained using the whole-cell patch-clamp technique. Micropipettes pulled from borosilicate glass were filled with an internal solution containing (in mM): 142 K-gluconate, 3 KCl, 10 HEPES, 0.5 EGTA, 5 phosphocreatine-tris, 5 phosphocreatine-Na2, 3 Mg-ATP, 0.5 GTP. Recording pipets had resistances of 2-5 MOhms.

Recordings occurred between 1.5 and 5 hours after the animal was sacrificed. Brain slices were transferred to a recording chamber and maintained at 34° C in oxygenated ACSF perfused at 2 mL/min. Electrophysiological recordings were restricted to layer 2/3 of the HVAs. These sites were identified using the fluorescence of the infected V1 axons in conjunction with anatomical landmarks to identify location along the anterior-posterior axis (**Figure 1D**). Online and *post-hoc* analysis ensured that we did not record from locations were cell bodies were infected with the optogenetic proteins. Thus, our measurements reflect the signals transmitted anterogradely from V1 to the higher areas. In all experiments, pyramidal cells were identified based on morphology and spiking responses. The majority of interneurons were identified with transgenic fluorescent labeling, but a small subset was assigned by somatic morphology and spiking properties (PV: 7/66; SOM: 2/62). For IN:Pyr EPSC ratios and L2/3 connectivity analysis we performed paired recordings of neighboring interneurons and pyramidal cells separated by fewer than 50 µm.

Neural signals were amplified using a MultiClamp 700B, low-pass filtered at 6 kHz, and digitized at 20 kHz using a Digidata 1550 (Axon Instruments). Data acquisition and stimulus presentation was controlled using the Clampex software package (pClamp 10.5, Axon Instruments). For characterization of intrinsic properties, we recorded in current clamp configuration and applied a series of current injections ranging from −680 pA to +680 pA. All neurons had <-55 mV resting membrane potential. For V1 axon stimulation experiments, optogenetically-evoked EPSCs were recorded in voltage clamp configuration while holding the membrane potential at the chloride reversal potential (−85 mV, uncorrected for liquid junction potential). Series resistance was monitored using −5 mV steps preceding each stimulus. We collected at least 10 sweeps for each recording. Only pairs where both neurons had <20 MOhms series resistance and stable holding current (<100 pA baseline variation) were included for analysis. Light pulses (350 µs) were generated using a 450 nm laser (Optoengine) coupled to the epifluorescence path (Olympus BX-RFA) and projected through a 40x water immersion lens (Olympus, 0.8 NA). The laser stimulation site had a diameter of 200 µm and was operated at a power of 20 mW/mm^2^ measured at the sample (**Figure 5A**). Light was targeted to layer 5 to avoid direct depolarization of the axon terminals.

For paired connectivity measurements between pyramidal cells and interneurons, we elicited a train of action potentials in one neuron while voltage clamping the other neuron to measure excitatory or inhibitory currents. The stimulus train consisted of ten 1 ms depolarizing steps at 12 Hz with a magnitude sufficient to reliably elicit an action potential on each step. EPSCs from pyramidal cells onto interneurons was measured by holding interneurons at −85 mV; IPSCs from interneurons onto pyramidal cells was measured by holding pyramidal cells at −40 mV. We recorded at least 10 sweeps for each direction.

### Anatomical analysis

For all post-hoc analysis of tissue, we manually selected regions of interest within each confocal z-stack. First, we aligned the image to be perpendicular to the surface of the brain; then we cropped the to include only the pia to the white matter, and to span 250 µm tangentially; finally, we only analyzed the subset of imaging frames from each confocal stack that exceeded half-maximal raw fluorescence to avoid including sections that were above or below the sample. Layer boundaries were assigned using size and density of cell bodies visualized with DAPI staining.

To quantify the density of axonal expression in the HVAs, we measured the profile of fluorescence intensity along the cortical depth. This was done by summing the raw fluorescence along the tangential dimension of the cortex. These depth-profiles were then aligned to the boundary between layer 1 and layer 2/3 and averaged across slices (4 per HVA per mouse). To compare axon density for all layers across HVAs, we first normalized the intensity to the maximum across areas, and then averaged across mice. This normalization serves to eliminate variability (e.g. injection amounts and incubation periods) across mice. To compare relative axon density distribution, we normalized to the maximum of the depth profile within each area, and then averaged across mice.

To quantify the density of interneurons in the HVAs, we developed a counting software written in MATLAB that allowed for the manual selection of fluorescent neurons. This allowed us to localize each neuron in the 3D volume of the coronal slice. Interneuron densities were calculated using the raw counts, and a tissue volume determined by the laminar boundaries, imaging window, and slice thickness. Interneuron densities were z-scored within layer and cell type. Distances between areas and layers were calculated as Euclidian distance between points using mean z-scores for PV and SOM densities, averaged across all mice.

### Electrophysiology analysis

Spike width, input resistance (R_in_), membrane time constant, spike frequency adaptation, and frequency-current relationship (FI) were calculated from current injection steps. Spike width was calculated as the full-width at half-max of the spike amplitude for the first spike at minimal depolarizing step intensity. Input resistance was calculated as the slope of the subthreshold V-I curve around rest. Membrane time constant was calculated by fitting a single exponential to the first 100 ms of membrane voltage change during the smallest hyperpolarizing step (−100 pA). I_h_ was calculated as the slope of the V-I curve between the two largest hyperpolarizing steps. Spike frequency adaptation was determined using the smallest depolarizing current injection sweep that elicited at least a 5 Hz initial firing rate, calculated as ISI_last_/ISI_first_. FI curves were generated by dividing total spikes during the current injection by the duration of the injection.

For analysis of optogenetically-evoked EPSCs recorded in simultaneously recorded interneurons and pyramidal cells, we averaged over all sweeps and smoothed the trial-averaged trace over 0.25 ms bins. IN:Pyr EPSC ratio, latency, rise time, and cumulative charge distributions were all calculated using this trial-averaged, smoothed trace. For calculating IN:Pyr EPSC ratios, we wanted to measure the maximum EPSC amplitude of V1 excitation and not polysynaptic connections from other pyramidal cells. Therefore, we used only the “early-onset” EPSC (EPSC_FF_) amplitude, which was identified using the first derivative of the trial-averaged current (*dI/dt*). The minimum of the derivative corresponds to the steepest rise of the of earliest excitatory input, presumably from action potentials elicited in V1 axons. This time point was then used to begin a 2 ms window over which we identified the maximum amplitude. The time of this first derivative minimum was determined individually for each neuron within pairs, such that time windows were often very similar but not identical (**Figure 5B**). The EPSC_FF_ maximum amplitude in each interneuron was divided by the paired pyramidal cell EPSC_FF_ maximum amplitude to generate the IN:Pyr EPSC ratio. Latency was calculated as time from laser onset to 20% of the EPSC_FF_ maximum amplitude. Rise time was calculated as time from 20-80% of EPSC_FF_ maximum amplitude. Jitter was calculated as the standard deviation of single trial latencies (time to 20% of individual trial EPSC_FF_ maximum amplitude).

To determine the relative distribution of excitatory charge in time, we created cumulative distributions of charge for all of the recorded EPSCs recorded. Using each cell’s calculated latency, we generated a cumulative distribution of charge over a 100 ms window beginning from response onset. The half-point was defined as the time to 50% of the total charge. Normalized traces used to compare currents across medial and lateral areas (**Figure 6E**) were generated by dividing the EPSC in each cell by its respective EPSC_FF_ maximum amplitude that was used to calculate IN:Pyr ratios.

For testing connectivity in each direction in our paired recordings, we averaged across all sweeps in the voltage clamped neuron. Current amplitudes were determined using the response to the first stimulus in the train. We categorized neurons as connected if there was an excitatory or inhibitory response to any of the stimuli in the train. Responses above baseline were determined with visual inspection but blinded to the area the pair was recorded in.

### Statistics

Unless otherwise stated all data are reported as mean ± SEM. Ratio data were log-transformed prior to statistical analysis. In other cases data were assumed to be normal. Unless otherwise stated, we used standard parametric tests (i.e. t-test and ANOVA with post-hoc Tukey test) and adjusted for multiple comparisons when necessary.

## Data availability

All anatomical and electrophysiological data supporting this manuscript are available on Mendeley Data (http://dx.doi.org/10.17632/rtnkp83yxs.1). Data were analyzed offline using custom MATLAB code. The code for data analysis is available on Github (https://github.com/Glickfeld-And-Hull-Laboratories/Manuscripts/tree/master/HVA_Inhibition).

## Acknowledgements

We thank Emily Burke, Courtney Dobrott, Kyra Leonard, and Wenjuan Kong for technical assistant with viral injections and *post hoc* histology; Kevin Murgas for technical assistance with cranial window surgeries and dye injections, and intrinsic imaging; Greg Horwitz and Yasmine El-Shamayleh for the gift of a virus encoding oChIEF; Court Hull and members of the Hull and Glickfeld laboratories for suggestions on experimental design and analysis; and Greg Field and Stephen Lisberger for comments on the manuscript.

## Author contributions

C.A.H., J.Y.L and L.L.G designed the experiments. I.M. and A.C.K. collected the anatomical data. C.A.H., I.M., A.C.K. and J.Y.L. analyzed the anatomical data. J.Y.L and C.A.H. collected and analyzed the electrophysiology data. J.Y.L. and L.L.G. wrote the manuscript with input from A.C.K.

## Figure Legends

**Figure 2 figure supplement 1.**
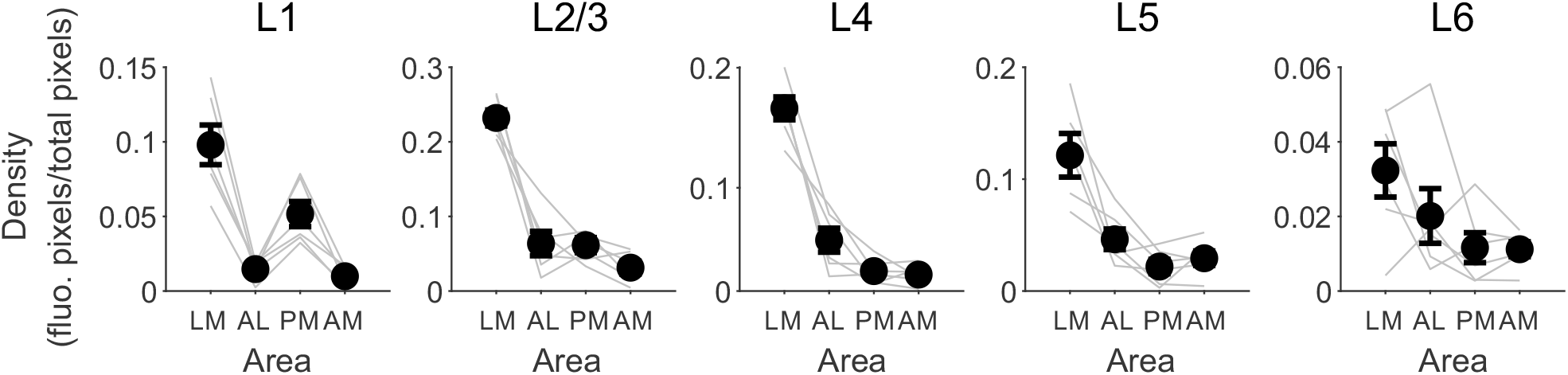
Non-normalized axon fluorescence density by area for L1-L6 (left to right). Grey lines are individual animals, black circles are mean across animals for each area and layer. Error bars are SEM across animals.

**Figure 3 figure supplement 1.**
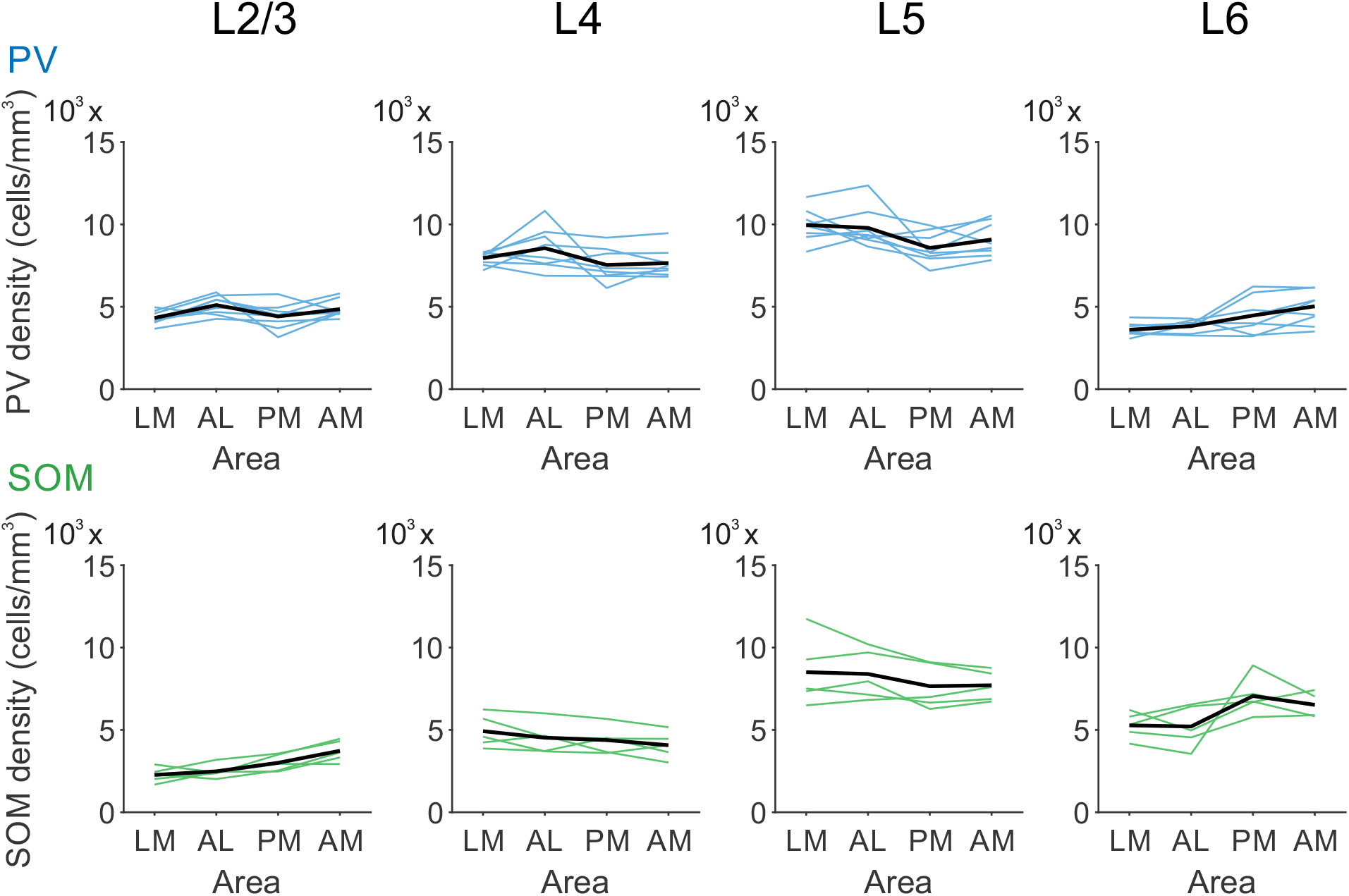
Top: Raw PV cell density by area for L2/3-L6 (left to right). Colored lines indicate individual animals, black line indicates mean across animals. Bottom: Same as top for SOM cells.

**Figure 6 figure supplement 1.**
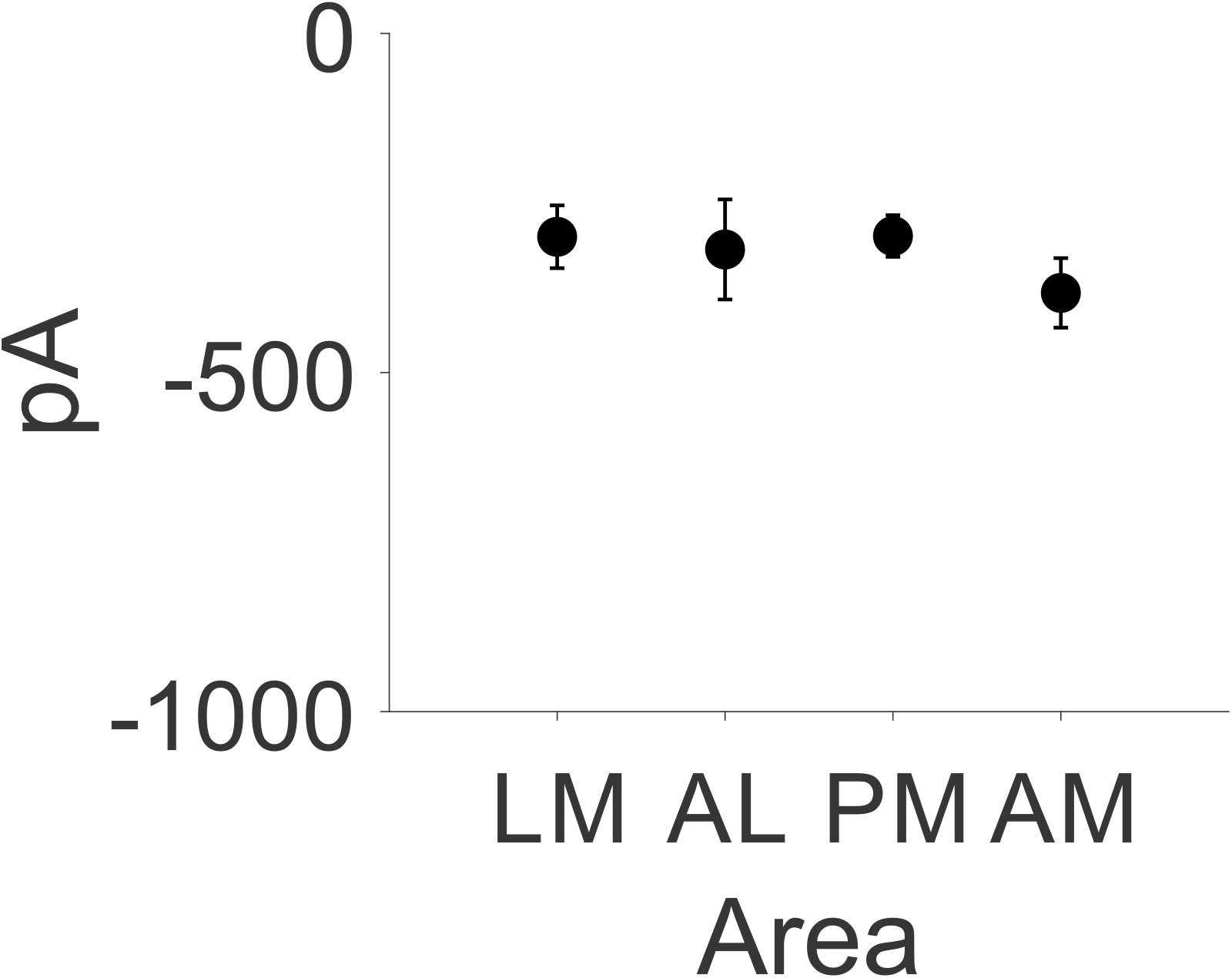
Raw pyramidal cell EPSC amplitude by area. Error bars are SEM across cells.

